# Elimination of Egfr-Overexpressing Cancer Cells by CD32 Chimeric Receptor T Cells in Combination with Cetuximab or Panitumumab

**DOI:** 10.1101/581017

**Authors:** Sara Caratelli, Roberto Arriga, Tommaso Sconocchia, Alessio Ottaviani, Giulia Lanzilli, Donatella Pastore, Carlo Cenciarelli, Adriano Venditti, Maria Ilaria Del Principe, Davide Lauro, Elisa Landoni, Hongwei Du, Barbara Savoldo, Soldano Ferrone, Gianpietro Dotti, Giuseppe Sconocchia

## Abstract

Cetuximab and panitumumab bind the human epidermal growth factor receptor (EGFR). While the chimeric cetuximab (IgG1) triggers antibody-dependent-cellular-cytotoxicity (ADCC) of EGFR positive target cells, panitumumab (a human IgG2) does not. The inability of panitumumab to trigger ADCC reflects a poor binding affinity of human IgG2 Fc for the FcγRIII (CD16) on NK cells. However, both human IgG1 and IgG2 bind the FcγRII (CD32) to a similar extent. Here, we have compared the ability of T cells, engineered with a novel low-affinity CD32^131R^ -chimeric receptor (CR), and those engineered with the low-affinity CD16^158F^–CR T cells in eliminating EGFR positive epithelial cancer cells (ECCs) in combination with cetuximab or panitumumab. Following T cell transduction, the percentage of CD32^131R^-CR T cells was (74±10) significantly higher than that of CD16^158F^-CR T cells (46±15). Only CD32^131R^-CR T cells bound panitumumab. CD32^131R^-CR T cells combined with the mAb 8.26 (anti-CD32) and CD16^158F^-CR T cells combined with the mAb 3g8 (anti-CD16) eliminated colorectal carcinoma (CRC), HCT116^FcγR+^ cells, in a reverse ADCC assay *in vitro*. Cross-linking of CD32^131R^-CR on T cells by cetuximab or panitumumab and CD16^158F^-CR T cells by cetuximab induced elimination of triple negative breast cancer (TNBC) MDA-MB-468 cells, and secretion of IFN gamma (IFNγ) and tumor necrosis factor alpha (TNFα). Neither cetuximab nor panitumumab induced Fcγ-CR T anti-tumor activity against KRAS-mutated HCT116, non-small-cell-lung-cancer, A549 and TNBC, MDA-MB-231 cells. ADCC of Fcγ-CR T cells was significantly associated with the over-expression of EGFR on ECCs. In conclusion, CD32^131R^-CR T cells are efficiently redirected by cetuximab or panitumumab against BC cells overexpressing EGFR.

**Article category:** Tumor Immunology and Microenvironment

**Novelty and Impact:** Monoclonal antibody-redirected Fcγ-CR T cell immunotherapy represents a promising approach in the fight against cancer. Here, we expand the application of this methodology to TNBC overexpressing the EGFR utilizing a novel CD32A^131R^-CR in combination with anti-EGFR mAbs. Our study supports the use of CD32A^131R^-CR T cells combined with panitumumab or cetuximab for targeting TNBC cells overexpressing the EGFR. Our results may be utilized as a platform for the rational design of therapies targeting TNBC overexpressing EGFR.

## INTRODUCTION

The lytic activity of ADCC is influenced by multiple variables including the type of affinity by which the Fc fragment of an antibody binds to the FcγR on a competent cytotoxic cell, the expression level of the targeted antigen on the surface of targeted cells, and the association constant of the antibody for the surface antigen of interest ^1^. The FcγR family includes CD16, CD32, and CD64. The former two receptors can be expressed in polymorphic forms, each of which displays different binding affinity for the Fc portion of IgG. The presence of valine at position 158 of CD16 (CD16^158V^) and of histidine at position 131 of CD32 (CD32^131H^) identifies the high-affinity receptors while the presence of phenylalanine at position 158 (CD16^158F^) and of arginine at position 131 (CD32^131R^) of CD16 and CD32, respectively, defines low-affinity receptors ^2^. Also, the activity of ADCC is influenced by the mAb subclasses since distinct subclasses have different association constants for the FcγRs CD16 and CD32. Cetuximab (IgG1) and panitumumab (IgG2) are currently utilized for the treatment of EGFR positive tumors. These two mAbs have demonstrated differential ability to mediate cell dependent cytotoxicity against EGFR positive epithelial cancer cells (ECCs) ^3^. Only cetuximab mediates CD16 positive NK cell-dependent cytotoxicity of EGFR positive cancer cells. The differential activity of cetuximab and panitumumab reflects the low affinity of IgG2 for the FcγR CD16 ^4^. In contrast, CD32 binds both IgG1 and IgG2 although with different affinity ^2^.

The role of ADCC in the *in vitro* and *in vivo* antitumor activity of tumor antigen (TA)-specific mAbs ^5^ has stimulated interest in genetically engineering T cells with the CD16 chimeric receptor (CD16-CR) ^6^. In these cells, the extracellular domain of CD16 was ligated to cytotoxic signaling molecules fused with ^7,8^ or without ^9,10^ T cell costimulatory molecules. This strategy allows the rapid generation of polyclonal T cells with a potent cytotoxic activity when combined with mAbs recognizing TAs expressed on tumor cell membrane. Based on this background information, we have utilized cetuximab and panitumumab as a model to demonstrate that CD32^131R^-CR T cells have higher cytotoxic activity than CD16^158F^-CR T cells since they eliminate EGFR positive cancer cells in combination with both cetuximab and panitumumab. In the present study, we have engineered T cells with a novel second generation of CD32^131R^-CR. The anti-tumor activity of CD32^131R^-CR T cells was compared to that of CD16^158F^-CR T cells in combination with cetuximab (IgG1) or panitumumab. Both engineered T cells, in combination with cetuximab, exerted a significant anti-tumor activity against breast cancer (BC) cells overexpressing the EGFR (EGFR^high^). However, only CD32^131R^-CR T cells effected cytotoxicity against EGFR^high^ BC cells in combination with either cetuximab or panitumumab. Our results strongly suggest that CD32^131R^-CR has a potential to be utilized in Fcγ-CR T cell-based immunotherapy of EGFR overexpressing BC cells.

## MATERIALS AND METHODS

### Antibodies and Reagents

Allophycocyanin (APC)-conjugated anti-human CD3 (cat. 555335), fluorescein isothiocyanate (FITC)-conjugated anti-human CD3 (cat. 555332), FITC-conjugated anti-human CD107A (cat. 555800), phycoerythrin (PE)-conjugated anti-human CD16 (cat. 555407), PE-conjugated anti-human CD32 (cat. 550586), FITC-conjugated mouse anti-human IgG (cat. 555786), FITC-conjugated goat anti-mouse IgG (cat. 555748), mouse anti-human CD3 (cat. 555329), and anti-human CD28 (cat. 555725) were purchased from BD Bioscience (San Jose, CA, USA). Mouse anti-human CD16 (clone 3g8), mouse anti-human CD247 (CD3ζ) mAb (clone 6B10.2) and mouse anti-human EGFR antibody (clone AY13) were purchased from Biolegend (San Diego, CA, USA). Anti-human CD32 mAb (clone 8.26) was purchased from BD Bioscience (San Diego, CA). Anti-human B7-H3 (CD276) mAb 376.96 was developed and characterized as described ^11^. mAb 376.96 was purified from ascitic fluid by affinity chromatography on Protein A. The activity and purity of mAb preparations was monitored by binding assays and SDS-PAGE. Cetuximab (Erbitux) and Panitumumab (Vectibix) were from Merck Serono (Darmstadt, Germany) and Amgen (Thousand Oaks, CA, USA), respectively. Anti-phospho Tyr142 (Y142) CD3ζ mAb (cat. ab68235) was purchased from Abcam (Cambridge, UK). 3-(4,5-Dimethylthiazol-2-Yl)-2,5-Diphenyltetrazolium Bromide (MTT) was obtained from Sigma-Aldrich (Saint Louis, MO, USA). FcR blocking reagent (BR) was purchased from Miltenyi (Bergisch Gladbach, Germany). GeneJuice® Transfection Reagent (Novagen) was from Millipore (Burlington, MA, USA). Human recombinant interleukin-7 (IL-7) and interleukin-15 (IL-15) were from Peprotech (London, UK). Lipofectamine 2000 was from Life Technologies (Carlsbad, CA, USA), and Retronectin (Recombinant Human Fibronectin) was purchased from Takara Bio (Saint-Germain-en-Laye, France).

### Cell lines

The 293T packaging cell line was used to generate the helper-free retroviruses for T cell transduction. 293T cells were cultured in Iscove’s Modified Dulbecco’s Medium (IMDM) supplemented with 10% Fetal Bovine Serum (FBS), 2mM L-glutamine, 0.1mg/mL streptomycin and 100U/ml penicillin hereafter referred to as IMDM complete medium (CM). KRAS-mutated A549 and HCT116 cell lines were maintained in RPMI-1640 CM. KRAS-mutated TNBC cells, MDA-MB-231, and KRAS wild-type TNBC cells, MDA-MB-468, were cultured in Dulbecco’s Modified Eagle’s Medium (DMEM) CM (Thermo Fisher Scientific, Waltham, MA, USA). 293T cells were kindly provided by Dr. Gianpietro Dotti, University of North Carolina, Chapell Hill, USA. A549 cells were kindly provided by Dr. Antonio Rossi, National Research Council, Italy. MDA-MB-231 and MDA-MB-468 cells were kindly provided by Dr. Maria Lucibello, National Research Council, Italy. HCT116 cells were kindly provided by Dr. Giulio Cesare Spagnoli, University of Basel, Switzerland. Mycoplasma-free cancer cell lines utilized in our study are part of our lab collection. Authentication test was successfully performed on November 21th, 2018 by PCR-single-locus-technology (Eurofins, Ebersberg, Germany). Cell lines were passaged for 4 to 8 times before use or kept in culture for a maximum of 6 weeks.

### CD32^131R^-CR construction

The signal peptide (nucleotide 1 - 101) and the extracellular region (nucleotide 102 - 651) of the low-affinity variant CD32A^131R^, hereafter referred to as CD32^131R^, was amplified by reverse-transcriptase polymerase chain reaction (RT-PCR) from RNA extracted from freshly isolated peripheral blood mononuclear cells (PBMCs) utilizing the following primers: forward 5’-GAGAATTCACCATGACTATGGAGACCCAAATG-3’ and reverse 5’-CGTACGCCCCATTGGTGAAGAGCTGCC-3’ (Thermo Fisher Scientific, Waltham, MA, USA). The PCR product was fused in tandem by restriction enzyme-compatible ends with the CD8α transmembrane domain and the CD28 and CD3ζ intracellular regions already available in the lab (CD32^131R^-CR). The generation of CD16^158F^-CR has already been described ^8^. The CD32^131R^-CR and CD16^158F^-CR genes were subcloned into the NcoI and MluI sites of the SFG retroviral vector.

### Retrovirus production and T cell transduction

Retroviral supernatants were obtained by transient transfection of 293T packaging cells, using the GeneJuice reagent, with the following vectors: the Peg-Pam vector containing the Moloney murine leukemia virus gag and pol genes, the RDF vector containing the RD114 envelope and the CD32^131R^-CR or CD16^158F^-CR SFG retroviral vectors. Forty-eight and 72h post-transfection, the conditioned medium containing the retrovirus was harvested, filtered, snap frozen, and stored at −80°C until use. For the generation of Fcγ-CR T cells, PBMCs (0.5×10^6^ PBMCs/ml) were cultured for 3 days in a non-tissue culture treated 24-well plate pre-coated with 1μg/ml of anti-CD3 and 1μg/ml of anti-CD28 mAbs in the presence of 10 ng/ml of IL-7 and 5ng/ml of IL-15. The viral supernatant was loaded on retronectin-coated non-tissue culture treated 24 well plates and spun for 1.5h at 2000xg. Activated T cells were seeded into the retrovirus loaded-plate, spun for 10’, and incubated for 72h at 37°C in 5% CO_2_. After transduction, T cells were expanded in RPMI-1640 CM supplemented with 10ng/ml of IL-7 and 5 ng/ml of IL-15 for 12-13 days and analyzed.

### Western blot

CD32^131R^-CR transduced and non-transduced T cells were lysed with Triton buffer composed of 1% (v/v) Triton X-100, 20mM Tris-HCL pH7.6, 137mM NaCl, 1mM MgCl_2_, 1mM CaCl_2_, 2mM phenylmethylsulfonyl fluoride (PMSF) supplemented with phosphatase (Sigma-Aldrich, Saint Louis, MO, USA) and protease (Roche, Basel, Switzerland) inhibitor cocktails. Thirty micrograms of protein lysates were resolved on Bolt 4-12% Bis-Tris plus gel (Invitrogen, Carlsbad, CA, USA) under reducing conditions and transferred to a nitrocellulose filter. The filter was probed overnight at 4°C with a mouse anti-human CD3ζ or rabbit anti-phospho-tyrosine CD3ζ (Y142) antibody. The latter was detected utilizing a horseradish peroxidase-conjugated donkey anti-mouse (Jackson Laboratory, Bar Harbor, ME, USA) for 1h at room temperature. Antibody binding was visualized with Amersham ECL Western blotting detection reagent (GE Healthcare, Little Chalfont, UK).

### Binding assay

A direct immunofluorescence analysis was utilized to test the Fc antibody-binding ability of a FITC-conjugated anti-CD107A mAb to the CD32^131R^-CR and CD16^158F^-CR. CD32^131R^-CR and CD16^158F^-CR T cells were incubated with 5μl of FITC-conjugated anti-CD107A, with or without Fc receptor blocking reagent (FcR BR) for 30 min at at 4°C. Then, cells were washed and analyzed by flow cytometry. Cetuximab or panitumumab Fc fragment binding to CD32^131R^-CR or CD16^158F^-CR T cells was evaluated by staining with a FITC-conjugated anti-human IgG.

### Flow cytometry

Expression of CD32^131R^-CR or CD16^158F^-CR on transduced T cells was assessed by staining for 30 min at 4°C with FITC-conjugated anti-human CD3, PE-conjugated anti-human CD32 or PE-conjugated anti-human CD16 mAbs, respectively. Cells were then analyzed by a 2-laser BD FACSCalibur (Becton Dickinson, Franklin Lakes, NJ, USA) flow cytometer. Results were analyzed utilizing Tree Star Inc. FlowJo software.

### Cytokine release assay

CD32^131R^-CR or CD16^158F^-CR transduced T cells (2×10^5^/well) were added to 96 well plates previously coated with 10μg/ml of anti-CD3, 3g8 or 8.26 mAbs. In co-culture experiments, CD16^158F^-CR T cells or CD32^131R^-CR T cells were plated in 96-well plates with target cell lines at 5:1 E:T ratio in the presence or absence of 3μg/ml of cetuximab or panitumumab or the anti-B7-H3 mAb, 376.96. Supernatants were collected after 24 or 48h of culture. IFNγ and TNFα levels were measured by ELISA (Thermo Fisher Scientific, Waltham, MA, USA).

### *In vitro* tumor cell viability assay

Tumor target cells (7×10^3^/well) were seeded into 96-well plates and CD16^158F^-CR T cells or CD32^131R^-CR T cells (35×10^3^/well) were added in the presence or absence cetuximab or panitumumab or the anti-B7-H3 mAb 376.96 (3μg/ml) (see above). Following a 48h incubation at 37°C, non-adherent T cells were removed. Then a suspension of fresh medium (100μl/well) supplemented with MTT (20μl/5mg/ml) was added to the adherent cells for 3h at 37°C. MTT was then removed and 100μl of dimethyl sulfoxide was added to each well. Absorbance (optical density, OD) was measured at 570 nm.

### Statistical analysis

Results were analyzed by a Paired-*T*-test or a Mann-Whitney test. The relationship between the two variables was measured by the Spearman’s rank correlation coefficient. Differences with *p*-value < 0.05 were considered significant.

## RESULTS

### CD32^131R^ and CD16^158F^ CRs are differentially expressed on T cells

Activated T cells were transduced *in vitro* with a gamma-retroviral vector encoding the CD32^131R^-CR (fig. 1A). Cells were then tested for expression of CD32^131R^-CR by western blot and flow cytometry analysis. For biochemical analysis, we utilized two mAbs specific for the non-phosphorylated and phosphorylated CD3ζ chain. Both mAbs detected 2 distinct bands. The band of a MW slightly higher than 51 kDa matches with the expected size of the CD32^131R^-CR while the smaller 18 kDa band detected in both control and CD32^131R^-CR expressing T cells corresponds to the endogenous CD3ζ chain (fig. 1B). By flow cytometry, CD32^131R^-CR was clearly detectable on the cell surface of engineered T cells (fig. 1C, left panel). Transduction efficiency of CD32^131R^-CR was significantly higher than that of CD16^158F^-CR (74% ± 10% vs. 46% ± 15%, p<0.001) (fig. 1C, right panel).

**Figure 1.**
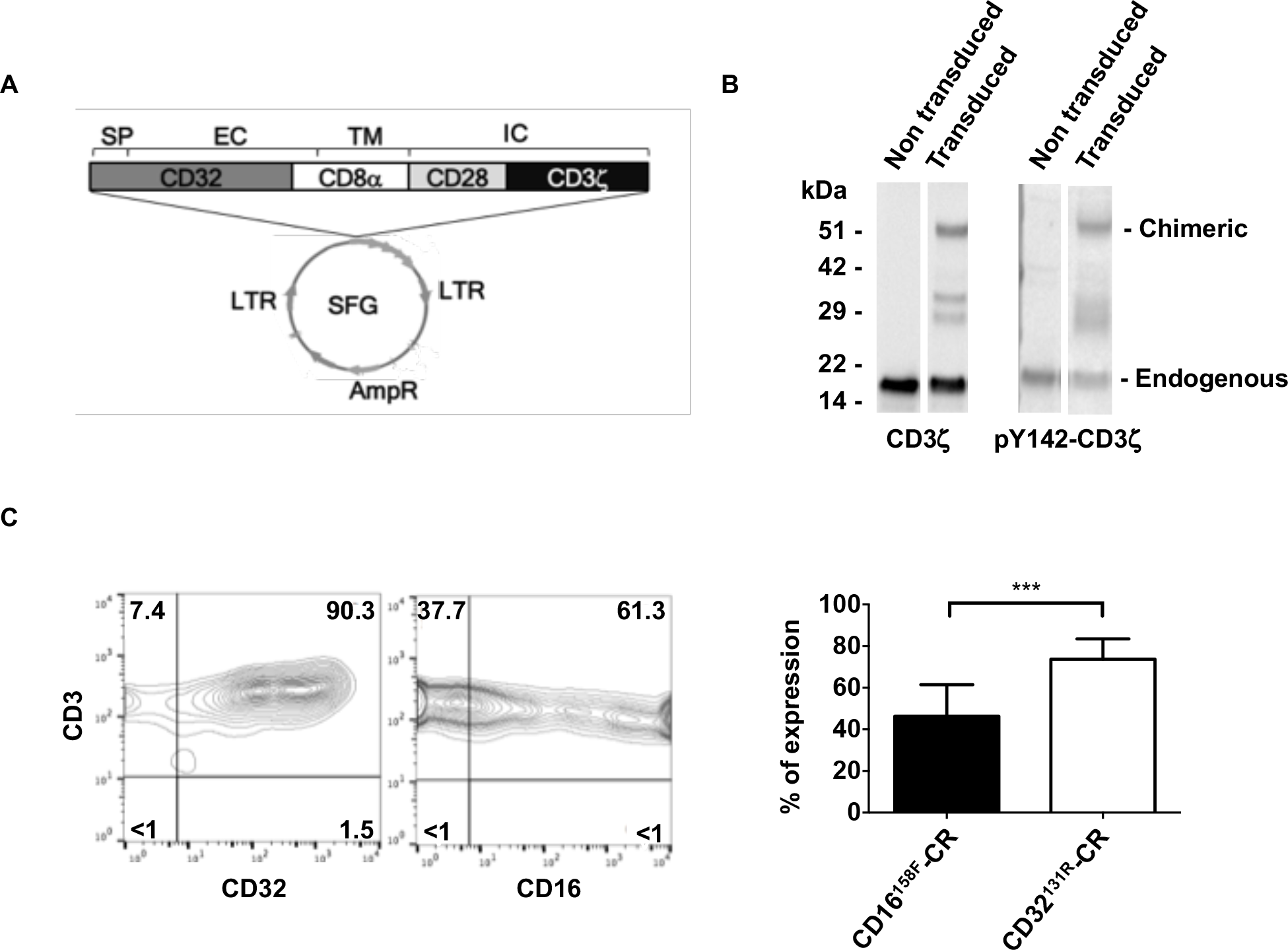
Molecular structure and expression efficiency of CD32^131R^-CR in T cells. A). Schematic representation of the CD32^131R^-CR gene cloned into the SFG retroviral vector. The CR transgene included the signal peptide (SP) and the extracellular domain (EC) of CD32^131R^ fused in tandem with the transmembrane (TM) region of CD8α and the intracellular (IC) signal motifs of CD28 and CD3ζ. B). Western blot analysis of CD3ζ expression in CD32^131R^-CR T cells. Soluble fractions of the cell lysates were separated, under reducing conditions, by SDS-PAGE gel electrophoresis and transferred on nitrocellulose filter membranes. Non-phosphorylated and phosphorylated CD3ζ were immunoblotted utilizing anti-CD3ζ and anti-phospho-Y142-CD3ζ mAbs, respectively. Lower bands refer to endogenous CD3ζ in both transduced and non-transduced T cells. Upper bands show the chimeric CD3ζ with the expected molecular weight specifically detected in transduced T cells. C). Following Fcγ-CR T cell transduction, cells were cultured for 3 days at 37°C in the presence of IL-7 and IL-15. Cells were then stained with FITC-anti-CD3 and PE-anti-CD32 or PE-anti-CD16 mAbs. Cells were analyzed by flow cytometry utilizing a BD-FACSCalibur^TM^. The frequency of Fcγ-CR^+^ T cells was calculated by evaluating percentages of CD3^+^CD32^+^ or CD3^+^CD16^+^ cells. This panel shows a representative experiment (left) and a cumulative analysis (right) of the results obtained from 10 different experiments. Numbers in the quadrants: % of cells; ***: p ≤ 0.001.

### CD32^131R^-CR specifically bound the Fc fragment of soluble immunoglobulins

In initial experiments, we compared the ability of CD32^131R^-CR and CD16^158F^-CR T cells to bind soluble IgG Fc fragment. As a model reagent, we chose the H4A3 mAb, a FITC-conjugated IgG1 specific for CD107A, an intracellular lysosomal-associated membrane protein (LAMP-1). We first evaluated the binding of anti-CD107A mAb on the surface of CD32^131R^-CR T cells in comparison to CD16^158F^-CR T cells. Following a 30 min incubation at room temperature, CD32^131R^-CR T cells effectively bound anti-CD107A mAb on their surfaces (fig. 2A, left panel) whereas CD16^158F^-CR did not (fig. 2A, right panel). Binding was highly specific since it was abrogated in the presence of FcR blocking reagent (BR). To further evaluate whether CD32^131R^-CR T cells were capable of binding mAb Fc fragment, in a more physiological condition, we tested whether the Fc fragment-binding capacity of CD32^131R^-CR was preserved in the presence of human immunoglobulins. Therefore, we incubated anti-CD107A with CD32^131R^-CR T cells in a buffer containing 10% of human plasma (fig. 2B). Following a 30 min incubation, at room temperature, anti-CD107A mAb was still bound to engineered T cells (fig. 2B).

**Figure 2.**
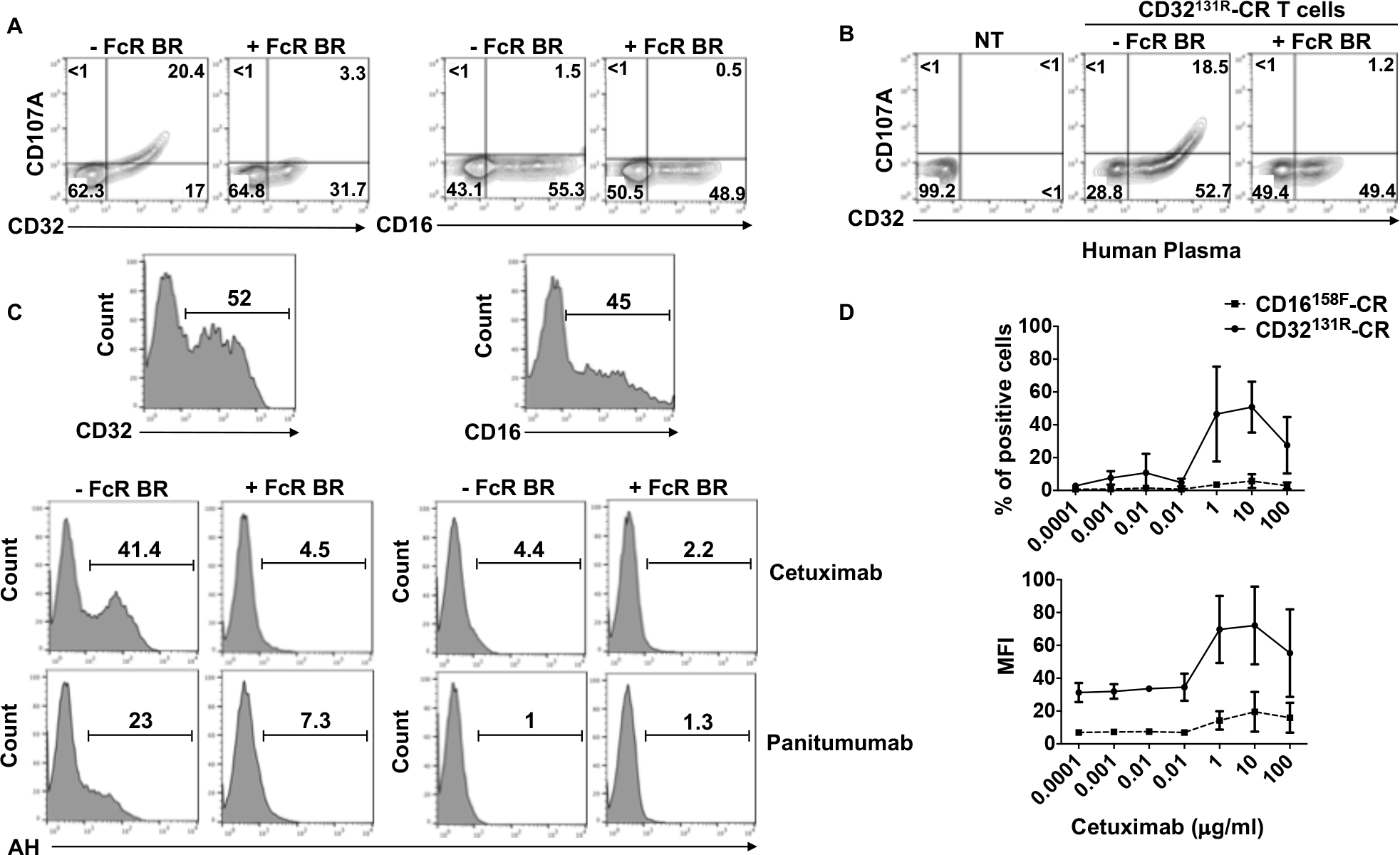
Analysis of the binding of mAb Fc fragments to CD32^131R^-CR and CD16^158F^-CR engineered T cells. The ability of CD32^131R^-CR T cells or CD16^158F^-CR T cells to bind specifically the Fc fragment of a FITC-conjugated anti-CD107A mAb was assessed. Following T cell transduction, blast cells were incubated for 20 min at 4°C with or without FcR BR in the absence (panel A) or presence (panel B) of 10% of human plasma. T cells were then washed and stained with fluorescent mAbs specific for the indicated markers and analyzed by flow cytometry. The binding of cetuximab or panitumumab Fc fragment on CD32^131R^-CR T cells (panel C, left) or CD16^158F^-CR T cells (panel C, right) was evaluated upon incubation with 3µg/ml of either cetuximab or panitumumab for 30 min at 4°C with or without FcR BR. Cells were then washed and stained with a FITC-conjugated mouse anti-human Ig antibody (AH). The binding of cetuximab and panitumumab was analyzed by flow cytometry. The maximum binding capacity of CD32^131R^- and CD16^158F^-CRs by cetuximab Fc fragment was determined in dose-response experiments (panel D). CD32^131R^-CR T cells (solid line) or CD16^158F^-CR T cells (dashed line) were incubated with increasing doses of cetuximab for 30 min, washed and then incubated with FITC-conjugated mouse anti-human Ig antibodies. Numbers in quadrants refer to percentages of positive cells. BR: blocking reagent, NT: not transduced.

To assess the Fcγ-CR T cell potential to target EGFR^+^ ECCs, upon incubation with anti-EGFR mAbs, we tested their antibody-binding capacity by utilizing cetuximab and panitumumab mAbs. Only CD32^131R^-CR T cells bound the Fc fragment of both soluble anti-EGFR mAbs (fig. 2C middle, panels) with higher binding capacity for cetuximab, as compared to panitumumab (fig. 2C, lower panels). Of note, binding of both mAbs was abolished in the presence of FcR BR (fig. 2C). Finally, we performed a dose-response binding assays, in which cetuximab was incubated at increasing concentrations with the Fcγ-CR T cells for 30 min at 4°C. As shown in Fig. 2D, only CD32^131R^-CR T cells (solid lines) bound cetuximab. The maximum binding capacity of CD32^131R^-CR, expressed as both percentages and MFI of positive cells, was achieved at concentrations ranging between 1-10 μg/ml of cetuximab. These results demonstrate that CD32^131R^-CR T cells have a superior binding ability for soluble mAbs than CD16^158F^-CR T cells.

### CD32^131R^-CR and CD16^158F^-CR T cells eliminate KRAS-mutated HCT116^FcγR+^ cells in redirected ADCC assays and release IFNγ and TNFα upon specific antigen stimulation

Next, we tested the ability of Fcγ-CR-transduced T cells to elicit cytotoxic activity in a reverse ADCC assay (fig. 3A). To this end, CD32^131R^-CR T cells and CD16^158F^-CR T cells were incubated in the presence of HCT116 cells, stably transfected with CD32, in the presence of the anti-CD32 and anti-CD16 mAbs respectively. Tumor cell viability was analyzed after 48h by MTT assay. Incubation with either CD32^131R^-CR T cells or CD16^158F^-CR T cells significantly reduced numbers of viable HCT116^FcγR+^ cells, consistent with efficient Fc-mediated cytotoxic activity. Furthermore, cross-linking of either Fcγ-receptor with specific mAbs induced the release of comparable amounts of IFNγ and TNFα (fig. 3B). These data indicate that both CD32^131R^-CR T cells and CD16^158F^-CR T cells clearly mediate comparable levels of reverse ADCC when given in combination with the mAbs 8.26 and 3g8 respectively.

**Figure 3.**
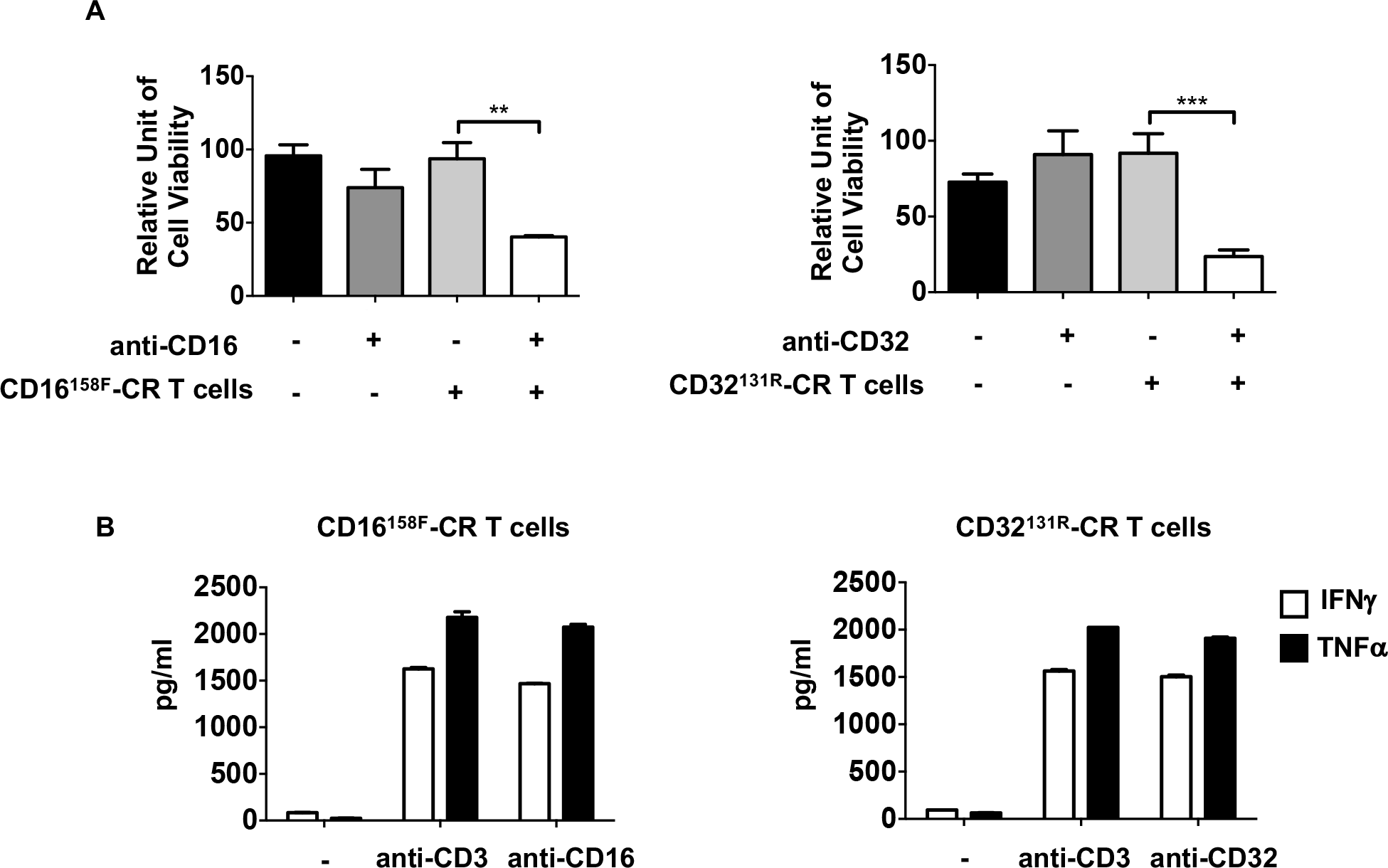
CD32^131R^-CR and CD16^158F^-CR engineered T cells are cytotoxic and produce equivalent amounts of IFNγ and TNFα. A) Redirected ADCC against HCT116^FcγR+^ cells. CD32^131R^-CR T cells or CD16^158F^-CR T cells were incubated with HCT116^FcγR+^ for 48h, at 37°C, at an E:T ratio of 5:1, with or without 8.26 (anti-CD32) or 3g8 (anti-CD16) mAbs. The viability of HCT116^FcγR+^ cells was evaluated by the MTT assay. B) Plastic bound 8.26 mAb or 3g8 mAb induces IFNγ and TNFα release in culture supernatants of Fcγ-CR T cells. Following cross-linking of CD32^131R^-CR or CD16^158F^-CR by plastic coated 8.26 or 3g8 as described in the method section, Fcγ-CR T cell supernatants were collected and IFNγ and TNFα contents were measured by ELISA.

### CD32^131R^-CR T cells and CD16^158F^-CR T cells differ in their ability to eliminate MD-MB-468 cells, in combination with cetuximab and panitumumab

The ability of CD32^131R^-CR to specifically bind the cetuximab Fc fragment prompted us to investigate whether this binding triggers ADCC against EGFR^+^ cancer cell lines. CD32^131R^-CR T cells and CD16^158F^-CR T cells were incubated with many ECC lines at an E:T ratio of 5:1. Tumor cell viability was assessed following a 48h incubation at 37°C. CD32^131R^-CR T cells significantly reduced the viability of MDA-MB-468 cells in the presence of cetuximab or panitumumab while CD16^158F^-CR T cells were only effective in the presence of cetuximab (fig. 4). In contrast, anti-B7-H3 379.96 mAb, which stained MDA-MB-468 cells, did not cause any detectable change in ECC viability. Neither cetuximab nor panitumumab had detrimental effects on MDA-MB-468 cells in the absence of Fcγ-CR T cells (fig. 4). However, CD32^131R^-CR T cells and CD16^158F^-CR T cells in combination with cetuximab or panitumumab failed to affect the viability of EGFR^+^ MDA-MB-231, A549, and HCT116 cells (fig. 4).

**Figure 4.**
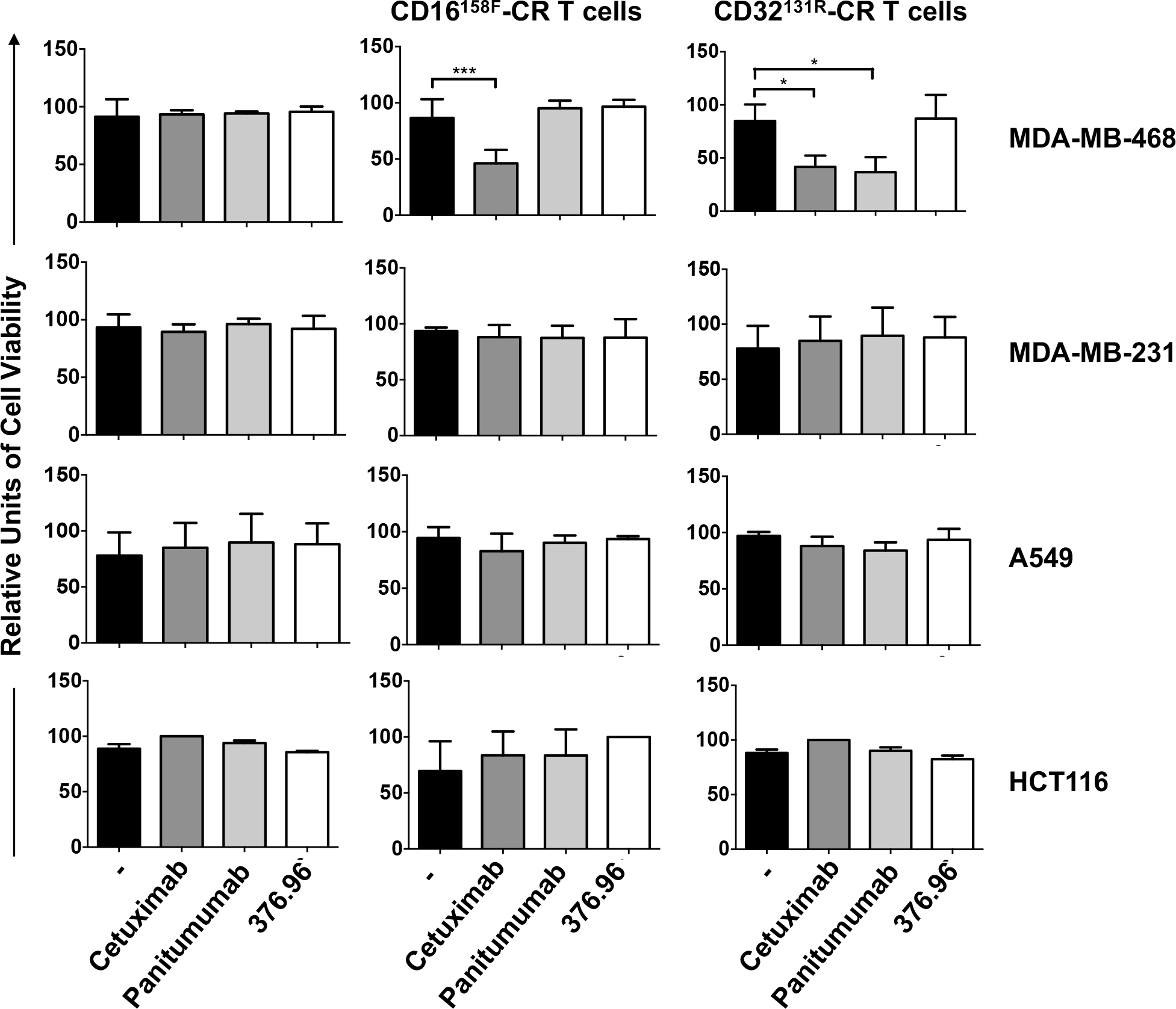
CD32^131R^-CR T cells eliminate MDA-MB-468 cells more efficiently than CD16^158F^-CR T cells. Anti-tumor activity of CD32^131R^-CR T cells and CD16^158F^-CR T cells was tested in combination with the indicated mAbs against EGFR^+^ and B7-H3^+^ cancer cells. TNBC cells, MDA-MB-468 and KRAS-mutated MDA-MB-231, NSCLC cells A549 and CRC cells HCT116, were chosen as target cells. MTT assays of tumor cell viability were performed following a 48h incubation, at 37°C, with CD32^131R^-CR or CD16^158F^-CR T cells with or without mAbs. The figure shows cumulative data, with mean ±SD values, of tumor cell viability obtained from 3 different donors at an E:T ratio of 5:1. Asterisks indicate: * = p<0.05 and *** = p<0.001.

Crosslinking of CD16^158F^-CR on engineered T lymphocytes cultured with MDA-MB-468 breast cancer cells promoted the release of IFNγ (fig. 5A) and TNFα (fig. 5B) in the presence of cetuximab but not of panitumumab. In contrast, both mAbs triggered the release of both cytokines by CD32^131R^-CR engineered T cells.

**Figure 5.**
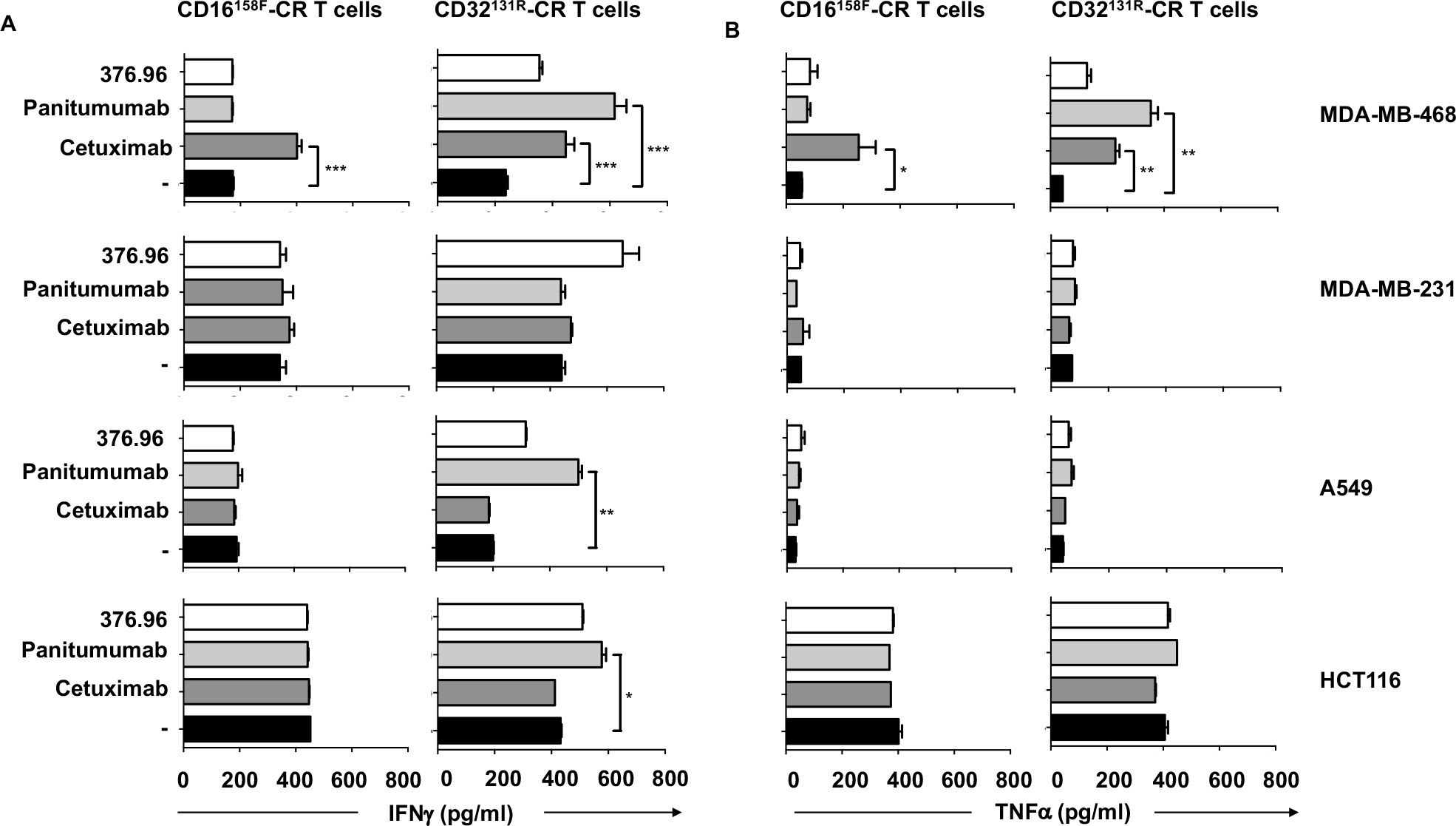
CD32^131R^-CR T cells secrete IFNγ and TNFα more efficiently than CD16^158F^-CR T cells following their conjugation with MDA-MB-468 cells by anti-EGFR mAbs. Following a 48h culture, at 37°C, of EGFR^+^ and B7-H3^+^ tumor target cells with CD16^158F^-CR T cells or CD32^131R^-CR T cells, in the presence or absence of the indicated mAbs, supernatants were harvested. The release of IFNγ (left) and TNFα (right) was measured in the culture supernatants by specific ELISA. The figure shows cumulative data, with mean ±SD values, of tumor cell viability from 3 different experiments performed at a 5:1 E:T ratio. Asterisks indicate: * = p<0.05, ** = p<0.01, and *** = p<0.001.

Furthermore, although panitumumab failed to mediate cell dependent cytotoxicity against the A549 and HCT116 cell lines, it induced a significant release of IFNγ by CD32^131R^-CR T cells (fig.5A) incubated with the two cell lines.

### Correlation of EGFR expression level on targeted cancer cell lines with the cetuximab dependent FcγCR T cell cytotoxicity

Our results indicate that among the EGFR^+^ cancer cell lines evaluated, only MDA-MB-468 cells were efficiently killed by CD16^158F^-CR T cells in combination with cetuximab and by CD32^131R^-CR T cells in combination with cetuximab or panitumumab (fig. 4). Since MDA-MB-468 cells express high levels of EGFR ^12^, we hypothesized that the ability of cetuximab to mediate ADCC activity of Fcγ-CR T cells against ECCs is associated with EGFR expression level on target cells. To test this hypothesis, we measured EGFR expression level on the surface of HCT116, A549, MDA-MB-231, and MDA-MB-468 cells and correlated it with the ability of cetuximab to mediate Fcγ-CR T cell cytotoxicity with target cells (fig. 6). As expected, MDA-MB-468 cells displayed the highest MFI upon staining with fluorochrome-labeled anti-EGFR mAb (fig. 6A). Furthermore, the ability of both CD16^158F^-CR and CD32^131R^-CR to reduce the viability of EGFR^+^ ECCs, in combination with cetuximab, displayed a highly significant correlation with the MFI of the target marker by the cancer cell lines tested (fig. 6B).

**Figure 6.**
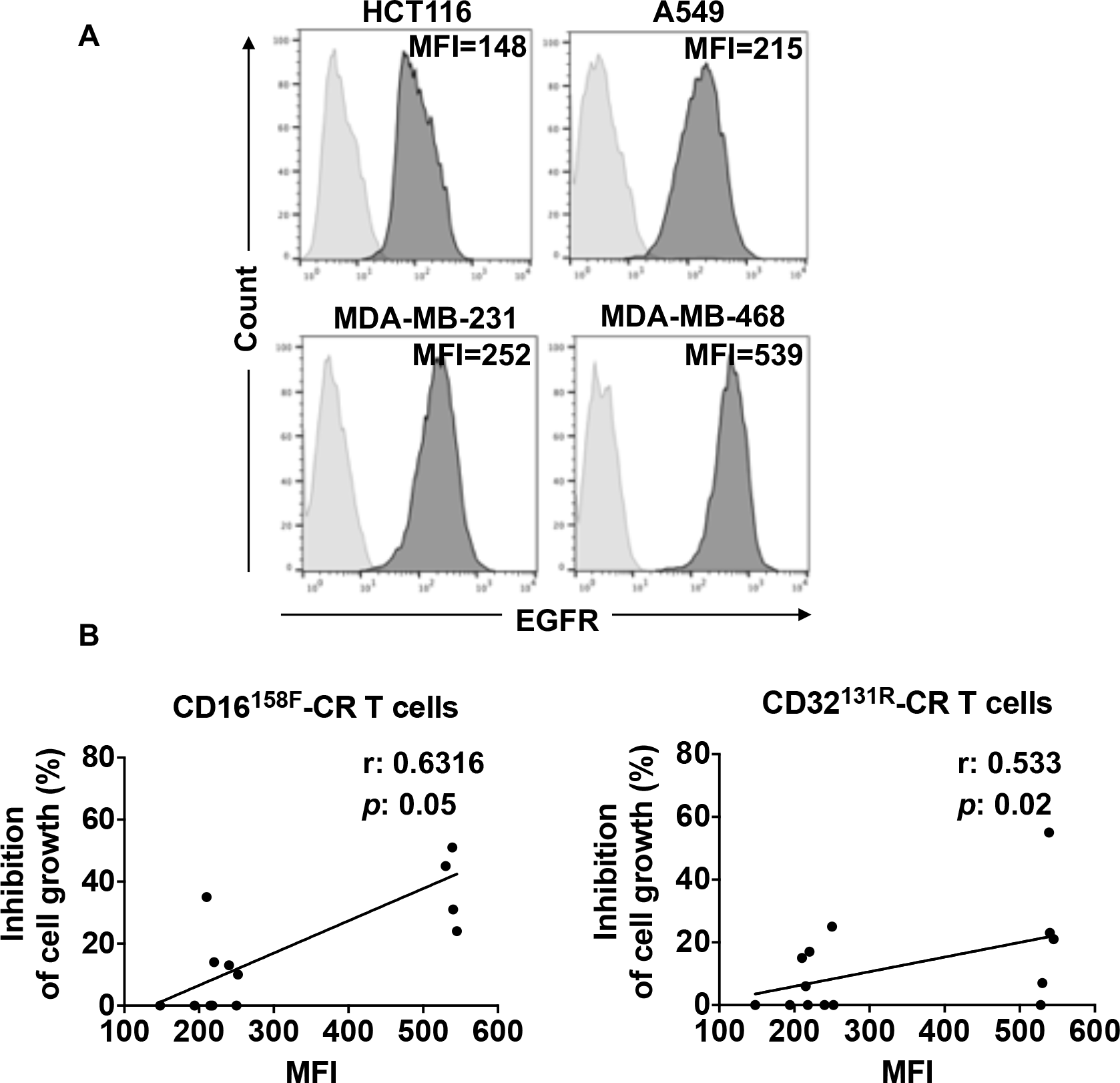
*In vitro* elimination of EGFR^+^ ECCs by CD32^131R^-CR and CD16^158F^-CR T cells is associated with EGFR overexpression. A) EGFR expression levels in tumor cell lines as assessed by flow cytometry. Cells were incubated with 3µg/ml of purified anti-EGFR antibody, washed and then stained with a FITC-conjugated anti-mouse IgG (gray filled histograms). Cells incubated with FITC-conjugated anti-mouse IgG were used as a negative control (light gray filled histograms). MFI is indicated for each cell line. B) Spearman’s correlation between EGFR expression levels (MFI) in HCT116, A549, MDA-MB-231, and MDA-MB-468 cell lines and the ECC elimination by CD16^158F^-CR T cells (left panel) and CD32^131R^-CR T cells (right panel) in combination with cetuximab. A regression line is reported in black. Spearman’s rank correlation coefficient (r) was 0.6316 for CD16^158F^-CR and 0.533 for CD32^131R^-CR.

## DISCUSSION

Rituximab and trastuzumab have been utilized to redirect first and second generation CD16^158V^-CR T cells against CD20^+^ and HER2^+^ hematologic and solid malignancies, respectively ^7,9,10^. The results obtained in pre-clinical studies suggest that CD16^158V^-CR T cells may act as universal chimeric receptor-effector cells capable of improving therapeutic effectiveness of TA-specific mAbs by an ADCC mechanism. The rationale, underlying the choice of the high-affinity extracellular CD16^158V^, for manufacturing Fc chimeras, is that CD16 triggers ADCC in NK cells ^13^. However, myeloid cells such as monocyte/macrophages and granulocytes can also mediate effector functions such as proinflammatory cytokine production ^14,15^ and cell-mediated cytotoxicity ^16^ including ADCC ^17,18^. Although CD16 is the major player in mediating ADCC, CD32 is also capable of promoting ADCC by myeloid cells ^17^. Similarly to CD16, CD32 is characterized by low (CD32^131R^) and high affinity (CD32^131H^) polymorphisms ^2^.

To date, there is scant information about the anti-tumor activity of CD16^158F^-CR T cells and the role of CD32^131R^-CR T cells is completely unknown. Growing experimental evidence suggests that CD16^158F^ and CD32^131R^ polymorphisms show differential binding affinities for IgG1 and IgG2 mAb Fc portions. Taking advantage from the availability of cetuximab (IgG1) and panitumumab (IgG2), here we demonstrate, for the first time, that both CD32^131R^- and CD16^158F^-CR T cells trigger ADCC to the TNBC cells, MDA-MB-468. In a side by side comparison, we show the superiority of CD32^131R^-CR over CD16^158F^-CR in redirecting engineered T cells against the MDA-MB-468 cells through anti-EGFR mAbs at least *in vitro*.

The transduction frequency of a retroviral vector encoding CD32^131R^-CR was significantly higher than that of CD16^158F^-CR. The higher expression frequency of CD32^131R^-CR on T cells may reflect the preferential propensity of hematopoietic cells to express CD32 as compared to CD16 ^19^. This result may be of relevance for manufacturing high numbers of selected engineered T cells for *in vivo* pre-clinical and clinical studies.

The affinity of CD16^158F^-CR and CD32^131R^-CR for IgG1 and IgG2 mAbs was significantly different. CD32^131R^-CR bound Fc fragments of soluble IgG1 (anti-CD107A and cetuximab) and to a lesser extent IgG2 (panitumumab) while CD16^158F^-CR T cells bound neither. The differential binding ability of cetuximab and panitumumab to CD32^131R^-CR is not surprising since CD32^131R^ polymorphisms bind IgG1 with a significantly higher affinity than IgG2. On the other hand, the failure of CD16^158F^-CR to bind soluble cetuximab (IgG1) is surprising since CD16^158F^ polymorphism has slightly lower ability to bind IgG1 than CD32^131R 2^. Our results are a bit different from those of Kudo et al. who showed a weak but detectable binding of soluble IgG1 mAbs, such as rituximab and trastuzumab, to CD16^158F^-CR. The different results obtained by Kudo et al ^7^ and by ourselves may reflect structural differences between the CD16^158F^-CR endodomain.

The induction of an effective ADCC of tumor cells by FcγR+ cytotoxic T cells needs to meet at least three distinct criteria. They include i) the presence of FcγR+ cytotoxic T cells with a functional lytic machinery; ii) FcγR binding affinity for the tested mAb Fc fragment sufficient to activate T cells; and iii) surface expression level of the antigen targeted by the tested mAb sufficient to activate effector mechanisms in T cells. Indeed, both CD16^158F^-CR and CD32^131R^-CR engineered T cells fully satisfy the first condition since anti-CD16 and anti-CD32 mAbs triggered a similar level of reverse ADCC when tested with KRAS-mutated, FcγR positive, HCT116 cells. However, CD16^158F^-CR T cells and CD32^131R^-CR T cells, in combination with cetuximab, neither released IFNγ and TNFγ and TNFα nor eliminated KRAS mutated ECCs including A549, HCT116, and MDA-MB-231 cells ^20,21^, although they were fully activated by the EGFR overexpressing MDA-MB-468 cells.

The differential antitumor activity of both Fcγ-CR T cells with wild type and KRAS-mutated ECC cells deserves some comments. The inability of both Fcγ-CR T cells to eliminate KRAS-mutated ECC cells does not reflect a mechanism of resistance of these target cells to the lytic activity of the two effector cells tested, since both of them can eliminate KRAS-mutated HCT116^FcγR+^ cancer cells in a redirect ADCC assay. On the other hand, the higher sensitivity of the wild type MDA-MB-468 than of the tested KRAS-mutated ECCs is likely to reflect the differential levels of surface EGFR expression. Indeed, the KRAS wild type, MDA-MB-468 cells overexpress EGFR ^22^ at a level higher than that on the KRAS-mutated ECCs utilized in this study. Our hypothesis is supported by Derer et al.’s finding that a KRAS mutation impairs the sensitivity of CRC cells to anti-EGFR mAbs because of C/EBPβ-dependent downregulation of EGFR expression ^23^.

The restoration of the sensitivity of KRAS-mutated ECCs to Fcγ-CR T cell lytic activity may require the generation of CD32-CR and CD16-CR with high affinity for the used mAb Fc fragments such as CD32^131H^-CR and CD16^158V^-CR. This strategy is supported by the ability of ECCs opsonized with anti-EGFR mAbs to induce a level of FCγR cross-linking insufficient to fully mediate ADCC, but sufficient to stimulate other FCγ-CR T cell functions such as cytokine production. In its support, we show that HCT116 and A549 cells, opsonized with panitumumab, promote IFNγ and TNFαrelease from CD32^131R^-CR T cells.

As a consequence, failure to generate an effective ADCC could be related to inadequate expression of EGFR on the surface of ECCs. In support of this hypothesis, CD16^158F^-CR T and CD32^131R^-CR T cells in combination with cetuximab damaged wild-type MDA-MB-468 cells overexpressing EGFR. Indeed, the results shown in figure 6 indicate that the extent of ECC reduced viability induced by the combination of either CD16^158F^-CR T cells or CD32^131R^-CR T cells with cetuximab directly correlated with EGFR expression levels on target ECCs. EGFR cross-linking MDA-MB-468 cells, with Fcγ-CR T cells and cetuximab, led to ADCC activation. The ability of CD16^158F^-CR T cells to mediate ADCC in the presence of cetuximab is somewhat unexpected since no binding of soluble cetuximab to these cells could be detected. This finding may reflect the ability of CD16^158F^-CR T cells to bind cetuximab only after EGFR cancer cell opsonization, which stabilizes ligand-receptor interactions.

Unlike cetuximab (IgG1), panitumumab (IgG2) did not induce significant ADCC by NK cells limiting its applications in cell-based cancer immunotherapy ^24^. However, panitumumab is still able to trigger ADCC by macrophages, which, with the exception of a small subset of cells ^25^, do not express CD16 but express CD32 and CD64 ^26^. As a logical consequence, engineering cytotoxic T cells with a CD32^131R^-CR has allowed us to demonstrate that panitumumab can stimulate strong ADCC by CD32^131R^-CR T cells against MDA-MB-468 cells overexpressing EGFR. These results open new perspectives for the use of panitumumab in cell-based targeted immunotherapy of solid tumors.

## Abbreviations

ADCC: antibody-dependent-cellular-cytotoxicity
APC: allophycocyanin
BC: breast cancer
CM: complete medium
CR: chimeric receptor
CRC: colorectal carcinoma
DMEM: Dulbecco’s Modified Eagle’s Medium
ECCs: EGFR positive epithelial cancer cells
EGFR: epidermal growth factor receptor
FBS: fetal bovine serum
FcR BR: Fc receptor blocking reagent
FITC: fluorescein isothiocyanate
IMDM: Iscove’s Modified Dulbecco’s Medium
IL-7: interleukin-7
IL-15: interleukin-15
INFγ: interferon gamma
mAb: monoclonal antibody
MFI: mean fluorescence intensity
NSCLC: non-small cell lung cancer
PBMCs: peripheral blood mononuclear cells
Pe: phycoerythrin
RT-PCR: reverse-transcriptase polymerase chain reaction
DMSO: dimethyl sulfoxide
OD: optical density
TA: tumor antigen
TNBC: triple negative breast cancer
TNFα: tumor necrosis factor alpha

## ACKNOWLEDGMENTS

This work was supported by the Italian Association for Cancer Research (AIRC) under grant IG17120. We thank Spagnoli G.C., Coccia M., and Rossi A. for technical support and Dr. Paggiolu M. and Dr. Papa P. for administrative assistance.

## Notes

**Disclosure of Potential Conflicts of Interest:** The authors declare no potential conflict of interest.

